# Shadow enhancers enable Hunchback bifunctionality in the Drosophila embryo

**DOI:** 10.1101/007922

**Authors:** Max V. Staller, Ben J. Vincent, Meghan D.J. Bragdon, Tara Lydiard-Martin Zeba Wunderlich, Javier Estrada, Angela H. DePace

## Abstract

Hunchback (Hb) is a bifunctional transcription factor that activates and represses distinct enhancers. Here, we investigate the hypothesis that Hb can activate and repress the same enhancer. Computational models predicted that Hb bifunctionally regulates the even-skipped (eve) stripe 3+7 enhancer (eve3+7) in Drosophila blastoderm embryos. We measured and modeled eve expression at cellular resolution under multiple genetic perturbations and found that the eve3+7 enhancer could not explain endogenous eve stripe 7 behavior. Instead, we found that eve stripe 7 is controlled by two enhancers: the canonical eve3+7 and a sequence encompassing the minimal eve stripe 2 enhancer (eve2+7). Hb bifunctionally regulates eve stripe 7, but it executes these two activities on different pieces of regulatory DNA–it activates the eve2+7 enhancer and represses the eve3+7 enhancer. These two “shadow enhancers” use different regulatory logic to create the same pattern.

**Significance statement:** Enhancers are regions of regulatory DNA that control gene expression and cell fate decisions during development. Enhancers compute the expression pattern of their target gene by reading the concentrations of input regulatory proteins. Many developmental genes contain multiple enhancers that control the same output pattern, but it is unclear if these enhancers all compute the pattern in the same way. We use measurements in single cells and computational models in *Drosophila* embryos to demonstrate that two enhancers that encode the same gene expression pattern compute differently: the same regulatory protein represses one enhancer and activates the other. Pairs of enhancers that output the same pattern by performing different computations may impart special properties to developmental systems.

## Introduction

Transcription factors (TFs) are typically categorized as activators or repressors, but many TFs can act bifunctionally by both activating and repressing target genes (1–4). Changes in TF activity can result from post translational modifications, protein cleavage or translocation of cofactors into the nucleus (5–7). However, in cases where a TF activates and represses genes in the same cells, bifunctionality is controlled by enhancer sequences, which are responsible for tissue-specific gene expression (8). For example, in *Drosophila*, Dorsal activates genes when it binds to enhancers alone or near Twist (9, 10) but represses genes when it binds near other TFs (11–13). TF binding site sequence can also alter TF activity, e.g. the glucocorticoid receptor (14, 15). Identifying how the activity of bifunctional TFs is controlled will be critical for inferring accurate gene regulatory networks from genomic data (16).

Here, we investigate how TF bifunctionality is controlled using a classic example: the *Drosophila* gene, *hunchback (hb)* (1, 20, 21). Hb both activates and represses *even-skipped (eve)* by acting on multiple enhancers. Hb activates *eve* stripes 1 and 2 and represses stripes 4, 5, and 6 (17, 18, 22, 39). Computational models from us and others support the hypothesis that Hb both activates and represses the enhancer that controls *eve* stripes 3 and 7 *(eve3+7)* (Fig. 1) (19, 23, 24).

**Figure 1.**
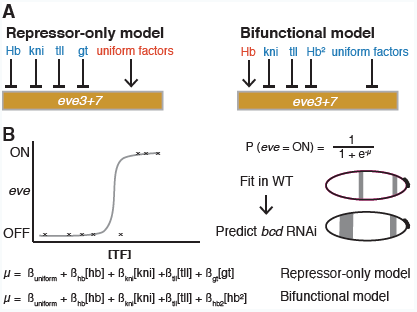
The repressor-only and bifunctional models formalize two alternative regulator sets for *eve* stripes 3 and 7. (A) The repressor-only model includes repression (red) by Hb, *knirps (kni), giant (gt),* and *tailless (tll)* and activation (blue) by a constant term that represents spatially uniform factors. The bifunctional model includes activation by a linear Hb term and repression by a quadratic Hb term, *kni, tll*, and uniform factors. (B) A schematic of the logistic regression framework. Logistic regression calculates the probability the target will be ON based on a linear combination of the concentrations of regulators (µ). We fit models in WT and use the perturbed regulator gene expression patterns to predict the perturbed *eve* patterns in *bcd* RNAi embryos.

In contrast to others, our computational models of *eve3+7* activity do not include regulatory DNA sequence (25–29). Instead, our modeling approach uses regression to identify activators and repressors that control a given pattern; we refer to the identity and role of the regulators as “regulatory logic.” Modeling regulatory logic without including DNA sequence enables a powerful strategy to dissect gene regulation in a complex locus. We can compare the regulatory logic of an enhancer reporter pattern to that of the corresponding portion of the endogenous pattern to determine if the annotated enhancer contains all relevant regulatory DNA.

Here we tested the hypothesis that Hb bifunctionally regulates *eve3+7*. We measured the endogenous *eve* expression pattern and that driven by an *eve3+7* enhancer reporter at cellular resolution under multiple genetic perturbations. We then used these data to challenge two computational models of *eve3+7* activity. In one model, Hb acts only as a repressor, while in the other, Hb acts as both an activator and a repressor (Fig. 1). The modeling indicated that *eve3+7* and the endogenous locus use different regulatory logic to position stripe 7. Specifically, *eve3+7* is only repressed by Hb, whereas the endogenous stripe 7 is both activated and repressed. We demonstrate that an additional sequence is activated by Hb and contributes to regulation of *eve* stripe 7 (17, 19, 27, 30–32). Thus, *eve* stripe 7 is controlled by a pair of shadow enhancers, separate sequences in a locus that drive overlapping spatiotemporal patterns (33). These shadow enhancers respond to Hb in opposite ways and use different regulatory logic.

## Results

### eve enhancer reporter patterns do not match the endogenous eve pattern

To determine if Hb bifunctionally regulates *eve3+7*, we compared the endogenous *eve* pattern to the pattern driven by a *lacZ* reporter construct in two genetic backgrounds (Fig. 2A). We refer to these data throughout the manuscript as “the *eve3+7* reporter pattern” and “the endogenous pattern.” We examined both wild-type (WT) embryos and embryos laid by females expressing short hairpin RNAs against *bicoid (bcd* RNAi embryos), where expression of all of the regulators, especially Hb, is perturbed (Figs S3, S4, 34). We quantitatively measured expression patterns at cellular resolution using *in situ* hybridization, 2-photon microscopy and an automated image processing toolkit (methods, 35, 36). We averaged data from many embryos into gene expression atlases (37). Importantly, the *eve3+7* reporter pattern results from the activity of *eve3+7* alone while the endogenous pattern integrates the whole locus.

**Figure 2.**
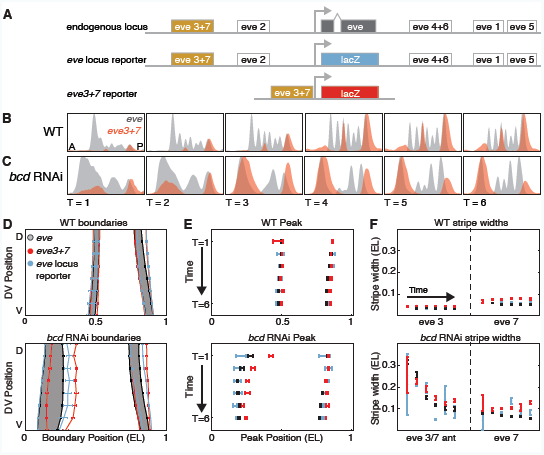
The *eve3+7* reporter pattern differs from the endogenous pattern. (A) The *eve* locus contains 5 annotated primary stripe enhancers. The endogenous pattern integrates the whole locus. The whole locus reporter pattern also integrates the whole locus, but the transcript is the same as the *eve3+7* reporter construct. The *eve3+7* reporter construct isolates the activity of the annotated enhancer sequence. (B) WT expression patterns are represented as line traces for a lateral strip of the embryo where anterior-posterior (A-P) position is plotted on the X-axis with expression level on the Y-axis. Endogenous *eve* pattern (gray), *eve3+7* reporter pattern (red). The reporter pattern was manually scaled to match the level of the endogenous pattern. (C) Line traces in *bcd* RNAi embryos. (D) The boundaries of the endogenous pattern (gray), the *eve3+7* reporter pattern (red), and the whole locus reporter pattern (blue) at T=3. All error bars are the standard error of the mean. The whole locus reporter pattern is more faithful to the endogenous pattern than the *eve3+7* reporter pattern, especially in the anterior of *bcd* RNAi embryos (eve 3/7 ant). The endogenous pattern is shaded for visual clarity. (E) Peak positions of stripes 3 and 7, calculated from the line traces in B and C. The *eve3+7* reporter pattern shows better agreement to the endogenous pattern in WT than in *bcd* RNAi embryos. (F) Stripe widths, calculated from the inflection point of the line traces in B and C. The *eve3+7* reporter pattern is wider than the corresponding endogenous pattern.

Our high resolution measurements revealed discrepancies between the endogenous pattern and the *eve3+7* reporter pattern. In WT embryos, the *eve3+7* reporter pattern overlaps the corresponding endogenous *eve* stripes, but these stripes are broader, have uneven levels, and the peaks lie posterior to the endogenous peaks (Fig. 2). These discrepancies were more pronounced in *bcd* RNAi embryos than in WT embryos, especially for the anterior stripe (Fig. 2D-F). When we tested reporters for other *eve* enhancers, we also found discrepancies between reporter patterns and the endogenous pattern (Figs S1, S2).

To test if the discrepancies between the *eve3+7* reporter pattern and the endogenous pattern resulted from differences in *eve* and *lacZ* transcripts, we measured the expression driven by a reporter encompassing the entire *eve* locus where the coding sequence had been replaced with *lacZ* (*eve* locus reporter, a generous gift from Miki Fujioka). In both WT and *bcd* RNAi embryos, the locus reporter pattern was more faithful to the endogenous pattern in terms of stripe peak positions and widths (Figs 2, S1, S2). Remaining differences between the endogenous and locus reporter patterns must arise from differences in the transcripts. Differences between the locus reporter and the *eve3+7* reporter patterns may arise from regulatory DNA outside of *eve3+7*. Together, these data suggest that the *eve3+7* reporter construct may not contain all the regulatory DNA that controls the expression of *eve* stripes 3 and 7.

### Different computational models capture the behavior of the endogenous locus and the enhancer reporter after Hb perturbation

We used computational models to dissect discrepancies between the *eve3+7* reporter pattern and the endogenous pattern. With our collaborators, we previously modeled the regulation of endogenous *eve* stripes 3 and 7 in WT embryos and simulated genetic perturbations that mimicked published experimental data (23). These models use logistic regression to directly relate the concentrations of input regulators to output expression in single cells. We constructed two models that together test the hypothesis that Hb can both activate and repress *eve* stripes 3 and 7. In the “repressor-only” model (the linear logistic model in Ilsley et al.), Hb has one parameter and only represses. In the “bifunctional” model (the quadratic logistic model in Ilsley et al.), Hb has two parameters that allow it to both activate and repress (Fig. 1). Both models performed equally well in WT embryos, but we favored the bifunctional model because it predicted the effect of a genetic perturbation. At that time, cellular resolution data for the *eve3+7* reporter pattern were not available, so we employed a standard assumption to interpret the models: the endogenous expression of *eve* stripes 3 and 7 could be attributed to the activity of the annotated *eve3+7* enhancer.

Here, we test this assumption explicitly by modeling the *eve3+7* reporter pattern and the endogenous pattern separately. Importantly, it is difficult to interpret the success or failure of a single model. It is much more powerful to compare the performance of two models that together formalize a hypothesis. We compared the performance of the repressor-only and bifunctional models in WT and *bcd* RNAi embryos. We used Hb protein and *giant* (*gt), tailless* (*tll*) and *knirps (kni)* mRNA as input regulators and thresholded the endogenous pattern and the reporter pattern for model fitting (Fig 1, methods). We report our modeling of the third timepoint, which is representative of results for other timepoints (Fig. S6), and evaluated model performance by computing the area under the receiver operating characteristic curve (AUC, 38).

We first analyzed the endogenous pattern: we fit our models in WT embryos and used the resulting parameters to predict expression in *bcd* RNAi embryos. Both models correctly predicted the positional shifts of stripe 7 and a wide anterior stripe, but the bifunctional model performed better than the repressor-only model (AUC_repressor_ = 0.93, AUC_bifunctional_ = 0.98, Figs 3F, S4).

**Figure 3.**
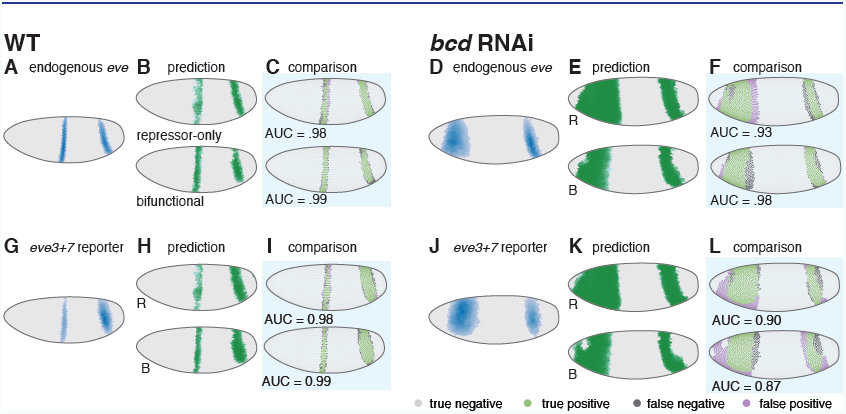
In *bcd* RNAi embryos, the bifunctional model more accurately predicts the endogenous pattern, and the repressor-only model more accurately predicts the *eve3+7* reporter pattern. (A) The endogenous *eve* pattern from in WT embryos shown as a rendering of a gene expression atlas. Cells with expression below an ON/OFF threshold (methods) are plotted in gray. For cells above this threshold, darker color indicates higher relative amounts. (B) The predictions of the repressor-only (top) and bifunctional (bottom) models in WT embryos. (C) Comparison of model predictions to the endogenous pattern in WT embryos. Green cells are true positives, purple cells are false positives, dark gray cells are false negatives, and light gray cells are true negatives. For visualization, the threshold is set to 80% sensitivity, but the AUC metric quantifies performance over all thresholds. (D) The endogenous *eve* pattern in *bcd* RNAi embryos. (E) The predictions of the repressor-only (top) and bifunctional models in *bcd* RNAi embryos. (F) Comparison of model predictions to the endogenous pattern in *bcd* RNAi embryos. The bifunctional model more accurately predicts the endogenous pattern in *bcd* RNAi embryos. (G-L) Same as A-F, respectively, for the *eve3+7* reporter pattern. The repressor-only model predicts the *eve3+7* reporter pattern more accurately in *bcd* RNAi embryos. Model parameters are in Table S1.

We next analyzed the *eve3+7* reporter pattern: again, we fit both models in WT embryos and used the resulting parameters to predict expression in *bcd* RNAi embryos. In this case, the repressor-only model was more accurate than the bifunctional model (AUC_repressor_ = 0.90, AUC_bifunctional_ = 0.87, Fig. 3L). We controlled for several factors that may confound prediction accuracy. We assessed sensitivity to changes in regulator concentrations, refit the models with *bcd* RNAi data, and refit the models on all of the data, none of which changed our conclusions (Figs S5 and S6, Supplemental Note 1).

These results suggest that Hb bifunctionally regulates the endogenous pattern but only represses the reporter pattern. Although the differences in relative model performance are subtle, the results support our hypothesis that the *eve3+7* reporter pattern is regulated differently than the endogenous pattern. However, these differences in model performance were not conclusive of their own accord and prompted us to return to the perturbation that previously distinguished the repressor and bifunctional models, ventral mis-expression of *hb* (23, 24).

### hb mis-expression confirms that the endogenous eve pattern and the eve3+7 reporter pattern respond to Hb differently

In Ilsley et al., we preferred the bifunctional model because it qualitatively predicted the behavior of a classic genetic perturbation. Mis-expressing *hb* along the ventral surface of the embryo (*sna::hb* embryos) causes *eve* stripe 3 to retreat and bend and stripe 7 to bend and bulge (39, Fig 4A and B). In simulations of this perturbation, the bifunctional model predicted this behavior while the repressor-only model predicted retreat of both stripes (Fig. 4 E and F reproduced from 23). We hypothesized that the endogenous and reporter patterns would respond differently to *hb* misexpression if Hb bifunctionally regulates the endogenous pattern but only represses the reporter pattern.

**Figure 4.**
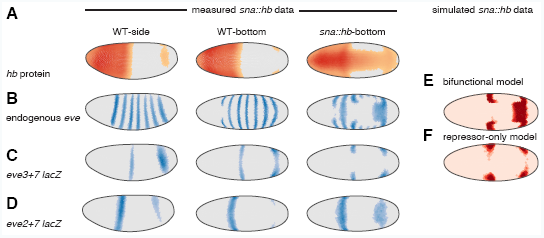
In hb ventral misexpression *(sna::hb)* embryos, the bifunctional model predicts the endogenous pattern while the repressor-only model predicts the *eve3+7* reporter pattern. (A) Hb protein in WT and *sna::hb* embryos (left, lateral view and right, ventral view). These data are computational renderings of gene expression atlases which average together data from multiple embryos (see Fig. S8 for number of embryos per time point). The relative expression level of each gene is shown in individual cells: cells with expression below an ON/OFF threshold (methods) are plotted in gray. For cells above this threshold, darker colors indicate higher levels. (B) Endogenous *eve* pattern. (C) The *eve3+7* reporter pattern. Both stripes retreat from ectopic Hb. (D) The *eve2+7* reporter pattern. Stripe 7 expands into the ectopic Hb domain. (E-F) Bottom (ventral) view of predictions of the bifunctional model (E) and repressor-only (F) models based on simulated *sna::hb* data. OFF cells are light pink and ON cells are red. Reproduced from Ilsley et al. 2013. All data and modeling from cohort 3 (other timepoints in Fig. S8).

We measured both patterns quantitatively at cellular resolution in *sna::hb* embryos (Fig. 4). As previously observed, the endogenous *eve* stripe 3 retreated from the ventral Hb domain and bent posteriorly, while the endogenous stripe 7 expanded and bent anteriorly, consistent with the bifunctional model (Fig. 4 B and E). By contrast, in the *eve3+7* reporter pattern, both stripes retreated from the ventral Hb domain, consistent with the repressor-only model (Fig. 4 C and F).

### Two shadow enhancers enable bifunctional Hb regulation of eve stripe 7

We hypothesized that additional regulatory DNA in the locus is activated by Hb to produce the *eve* stripe 7 bulge in *sna::hb* embryos. We tested an extended version of the minimal *eve2* enhancer for this activity based on several previous observations. Hb is known to activate the *eve2* enhancer (17, 18, 40, 42); longer versions of *eve2* drive stripe 7 in some embryos (27, 30, 31, 40); orthologous *eve2* enhancers from other species sometimes drive stripe 7 expression (32, 41); and finally, in *sna::hb* embryos, the border of the expanded stripe 7 appears to be set by *Krüppel (Kr)*, a known regulator of *eve2* (Fig. S7, 17, 42). The fragment we chose drives both stripes 2 and 7 (Fig. S8, Table S2); we call this enhancer reporter construct *eve2+7*.

In *sna::hb* embryos, the stripe 7 region of the *eve2+7* reporter pattern expanded, recapitulating the bulge observed in the endogenous *eve* pattern (Fig. 4B and D). We conclude that Hb activates endogenous *eve* stripe 7 through the *eve2+7* enhancer. Taken together, our results indicate *eve* stripe 7 expression is controlled by at least two enhancers with different regulatory logic.

## Discussion

To test whether Hb can activate and repress the same enhancer, we used quantitative data to challenge two computational models that formalize different roles for Hb. We measured expression of endogenous *eve* and transgenic reporter constructs at cellular resolution under two genetic perturbations. By comparing the regulatory logic of the endogenous and *eve3+7* reporter patterns, we uncovered two enhancers that both direct expression of *eve* stripe 7. These shadow enhancers direct the same pattern in different ways: one is activated by Hb while the other is repressed. This form of regulatory redundancy enables Hb to “drive with the brakes on” to control *eve* stripe 7.

### Two shadow enhancers control eve stripe 7 expression

Early studies suggested control of *eve* stripe 7 expression was distributed over DNA encompassing both the minimal *eve3+7* and *eve2* enhancers (17, 19, 30, 31, 40). We find that there are at least two pieces of regulatory DNA in this region that position stripe 7. The minimal *eve3+7* enhancer is repressed by Hb (19, 39, 43), while the *eve2+7* enhancer, which encompasses the minimal *eve2* enhancer, is activated by Hb. This activation may be direct or indirect. Based on the results presented here, we cannot rule out the possibility that the bulge of *eve2+7* in *sna::hb* embryos is indirect, due to activation by other TFs and retreat of Gt and Kni (39, 45). However, we hypothesize that Hb activation of *eve2+7* is direct. If Hb activation of *eve2+7* is indirect, Hb binding to *eve2+7* in these cells would have to have little or no effect on stripe 7 expression (44). Moreover, Hb binds to and activates the minimal *eve2* enhancer (17, 18, 40, 44, 69).

In addition to responding to Hb in opposite ways, the *eve2+7* and *eve3+7* enhancers are likely differentially sensitive to additional TFs. *eve3+7* is activated by Stat92E and Zelda (43, 46). The anterior border of stripe 7 is set by Kni repression, and the posterior border is set by Hb repression (19, 39, 43). The minimal *eve2* enhancer is activated by Bcd and Hb, its anterior boundary is set by Gt, and its posterior boundary set by Kr (17, 18, 40, 42). In agreement with others, we speculate that the anterior boundary of *eve* stripe 7 in *eve2+7* may be set by Gt (27). Taken together, this evidence argues that *eve3+7* and *eve2+7* position stripe 7 using different regulatory logic.

The molecular mechanism by which Hb represses and activates remains unclear. One hypothesis is that other TFs bound nearby convert Hb from a repressor into an activator as is the case for Dorsal (9, 12, 13). There is genetic evidence for activator synergy between Bcd and Hb (17, 47) and activator synergy between Hb and Caudal has been proposed by computational work (29). Another hypothesis is that Hb monomers are activators, but DNA-bound Hb dimers are repressors (24, 48). Testing these hypotheses will require quantitative data in additional genetic backgrounds and mutagenesis of individual binding sites in the two enhancers.

### Comparing the regulatory logic of reporter and endogenous patterns may be helpful for mapping regulatory DNA

“Veteran enhancer-bashers, and those who carefully read the papers, know that ‘minimal’ enhancer fragments do not always perfectly replicate the precise spatial boundaries of expression of the native gene…” (33). Our data clearly support this often neglected aspect of enhancer reporter constructs. One explanation offered for such discrepancies is different transcript properties. We controlled for this possibility and conclude that transcript properties contribute to the differences between reporter and endogenous patterns, but are not the only source. Here, we find that additional regulatory DNA in the locus also plays a role.

Finding all of the active regulatory DNA in a locus is challenging. Enhancer reporter constructs are powerful, but can only determine whether a piece of DNA is sufficient to drive a particular pattern in isolation when placed next to the promoter. By comparing the regulatory logic of the *eve3+7* reporter pattern and the endogenous pattern, we found a new feature of *eve* regulation. However, *eve3+7* and *eve2+7* may not contain all of the DNA that contributes to stripe 7 expression *in vivo.* Emerging technologies for manipulating the endogenous locus and larger reporter constructs will be helpful for comprehensively mapping regulatory DNA (49–51).

### The bifunctional model is a superposition of two computations

Models are not ends, but merely means to formalize assumptions and develop falsifiable hypotheses (52, 53). The bifunctional model accurately predicts the behavior of the endogenous *eve* stripe 7 pattern in WT and perturbed embryos, but it does not predict the behavior of either *eve3+7* or *eve2+7*. The interpretation in Ilsley et al. that Hb bifunctionally regulated *eve3+7* was based on a common assumption: that the endogenous pattern could be attributed to the annotated enhancer. Here we show that Hb bifunctionality is due to separate enhancers. Ilsley et al. interpreted the success of the bifunctional model as evidence for concentration-dependent control of Hb activation and repression, as has been proposed for Hb and other TFs (20, 24, 54). This interpretation cannot be true because Hb activates and represses in the same cells. Our favored hypothesis is that Hb bifunctionality is controlled by sequence features in each enhancer.

The bifunctional model effectively behaves as a superposition of the *eve3+7* and *eve2+7* enhancer activities to accurately predict the behavior of the endogenous locus. It is currently unclear how multiple active enhancers impinge on the same promoter, which makes it challenging to predict their combined behavior. The promoter may integrate information from multiple enhancers in various ways, ranging from independent addition to dominance of one enhancer due to a long-range repressor (33, 55, 56). The behavior of stripe 7 is not consistent with dominant repression by Hb, but we cannot rule out any other mechanisms. Elucidating how promoters integrate information will be critical for predicting the behavior of complex developmental loci where shadow enhancers are prevalent.

## Conclusion

By combining computational modeling and directed experiments, we uncovered a new feature of a highly-studied locus, long held up as a textbook example of enhancer modularity (57). We tested the hypothesis that Hb bifunctionally regulates the *eve3+7* enhancer and discovered that bifunctionality is due to two enhancers that respond to Hb in opposite ways. This example provides an opportunity to uncover how Hb bifunctionality is controlled, which will improve our ability to interpret regulatory DNA and infer connections in gene regulatory networks.

Regulatory redundancy in control of *eve* stripe 7 expression may have functional consequences. Shadow enhancers may arise from genetic drift (58); however, shadow enhancers in other developmental loci confer robustness to genetic or environmental stresses (55, 59, 60), facilitate temporal refinement of patterns (61), and/or increase expression synchrony and precision (62). This example demonstrates that shadow enhancers can use different regulatory logic to position the same pattern, which may have useful properties for the embryo.

## Materials and Methods

### Fly Work

The *bcd* RNAi gene expression atlas is described in Staller et al. 2014 (35) and available at depace.med.harvard.edu. Briefly, we combined short hairpin RNA knockdown of *bcd* with *in situ* hybridization and 2-photon imaging and automated image segmentation (37, 63–65). Hb protein stains used a guinea pig anti-*hb* from John Reinitz (University of Chicago, IL). Embryos were partitioned into six time points using the degree of membrane invagination (0-3%, 4-8%, 9-25%, 26-50%, 51-75%, and 76-100%) which evenly divide the ~60 min blastoderm stage (36). All enhancer reporters are in pBOY and integrated at attP2 (32, 66) (Table S2). The *eve* locus *lacZ* reporter was a gift from Miki Fujioka (Thomas Jefferson University, PA). *hb* ventral misexpression was performed as described in Clyde et al., 2003 using two copies of the *sna::hb* transgene on chromosome 2.

### Building the coarsely aligned sna::hb gene expression atlas

We determined the genotype of the *sna::hb* embryos by examining the *eve* or *fushi-tarazu (ftz)* mRNA patterns. Embryos were aligned morphologically to create a coarsely registered gene expression atlas (37). Data is available at depace.med.harvard.edu.

### Logistic modeling of enhancer gene regulatory functions

The logistic modeling framework was developed and described in detail previously (23). All modeling was performed in MatLab (MathWorks, Natick, MA) using the DIP image toolbox (diplib.org) and the PointCloudToolBox (bdtnp.lbl.gov). Ilsley et al. used protein data for Gt, whereas we used mRNA data. For genes where we used mRNA data, the mRNA and protein patterns are correlated (37, 67). For the enhancer *lacZ* reporters, we thresholded cells to be ON or OFF by creating a histogram of the expression data (50 bins), identifying the bin with the most counts and adding one standard deviation. Our ON set included all cells expressing the reporter, and our OFF set includes all other cells. All regulators are maintained as continuous values.

To threshold the endogenous WT *eve* pattern into ON and OFF cells we used 0.2 for all time points (23). To threshold the endogenous *eve* patterns in the *bcd* RNAi atlas, we used the lowest threshold that would separate the stripes: 0.1, 0.15, 0.15, 0.2, and 0.21 for T=2 through T=6 respectively. To compare the modeling of the reporter and the endogenous patterns, the ON set included all cells in the endogenous *eve* stripes 3 and 7 and the OFF set included all other cells. This OFF set is different from Ilsley et al., but this change does not have a large effect on the AUC scores in *bcd* RNAi embryos (Table S1).

### Sensitivity analysis

For the sensitivity analysis (Fig. S5), for each TF, we scaled the concentration of the *bcd* RNAi atlas *in silico* and recomputed the model AUC scores.

### Binding site predictions

For the *Kr* binding site analysis in Fig. S7, we predicted binding sites using PATSER (stormo.wustl.edu) with a position weight matrix derived from bacterial 1-hybrid data (68). Binding sites were visualized using InSite (cs.utah.edu/~miriah/projects).

### Quantifying concordance between reporters and endogenous patterns

For each embryo, we used the pointcloud toolbox in Matlab to find pattern boundaries by creating 16 anterior-posterior line traces and finding the inflection point of each trace. Finding the boundary by using half the maximum value of the stripe peak identifies a very similar boundary to the inflection point. To find the peaks of the endogenous and reporter stripes, we took one line trace along the lateral part of the embryo and found the local maxima.

## Acknowledgements

We thank Miki Fujioka for sharing the *eve* locus reporter flies ahead of publication, Tara Lydiard-Martin for making the enhancer reporter fly lines, and Steve Small for the *sna::hb* flies; Garth Ilsley for developing the initial models, help with figures and stimulating discussions; John Reinitz for the Hb antibody; Steve Small, Becky Ward, Garth Ilsley, Peter Combs, Alistair Boettiger, two anonymous reviewers and members of the DePace lab for comments on the manuscript. This work was supported by the Harvard Herchel Smith Graduate Student Fellowship (MVS), Jane Coffin Childs Memorial Fund for Medical Research (ZBW), NIH K99HD073191 (ZBW), and NIH U01 GM103804-01A1 (AHD).

